# Gα_s_-specific structural elements attenuate interactions with RGS proteins

**DOI:** 10.1101/2024.07.27.605439

**Authors:** Sabreen Higazy-Mreih, Meirav Avital-Shacham, Christian LeGouill, Michel Bouvier, Mickey Kosloff

## Abstract

Heterotrimeric (αβγ) G proteins are molecular switches that are activated by G protein-coupled receptors (GPCRs) and regulate numerous intracellular signaling cascades. Most active Gα subunits are inactivated by Regulators of G protein Signaling (RGS) proteins, which determine the duration of G protein-mediated signaling by accelerating the catalytic turn-off of the Gα subunit. However, the G protein Gα_s_ does not interact with known RGS proteins. To understand the molecular basis for this divergent phenomenon, we combined a comparative structural analysis of experimental and modelled structures with functional biochemical assays. This analysis showed that Gα_s_ contains unique structural elements in both the helical and the GTPase domains. Modeling identified helical domain insertions, missing in experimental structures, that project towards the interface with RGS proteins, and residues in the GTPase domain that might interfere with RGS binding. Mutagenesis of Gα_s_ and measurements of RGS GAP activity showed that three residues in the Gα_s_ GTPase domain are both necessary and sufficient to prevent Gα_s_ inactivation by RGSs. Indeed, substitution of all three Gα_s_ residues with the corresponding residues from Gα_i1_ enabled efficient inactivation by RGS proteins. These results shed new light on the mechanistic bases for G protein specificity towards RGS proteins.

## Introduction

Heterotrimeric (αβγ) G proteins are ubiquitous molecular switches that are activated by G protein coupled receptors (GPCRs) and control numerous downstream effectors and physiological systems (Gilman 1995, Sprang 1997, Oldham and Hamm 2008). In humans, 16 genes encode for Gα subunits that are classified into four subfamilies, based on Gα sequence similarity and the effectors they activate – the G_s_, G_i_, G_q_, and G_12/13_ subfamilies (Sprang 1997, Milligan and Kostenis 2006). The activation state of G proteins is regulated by the identity of the guanine nucleotides bound to the Gα subunit (Gilman 1995, Sprang 1997, Oldham and Hamm 2008), whereby Gα subunits are turned “on” by exchange of bound GDP for GTP and turned “off” by the hydrolysis of the bound GTP to GDP. While most Gα subunits are inactivated by regulators of G protein signaling (RGS) proteins that accelerate GTP hudrolysis, RGS proteins do not inactivate members of the G_s_ subfamily (Berman, Kozasa et al. 1996, Hunt, Fields et al. 1996, Natochin and Artemyev 1998, Masuho, Balaji et al. 2020). Yet, the molecular and structural determinants underlying this unique property of the G_s_ subfamily have not been fully elucidated.

Members of the G_s_ subfamily are expressed in almost all cells in mammals and include Gα_s_ and Gα_olf_, which are 88% identical in sequence (Jones, Masters et al. 1990), with almost all studies using Gα_s_ as the representative member of this subfamily (Sprang, Chen et al. 2007).

Members of the G_s_ subfamily stimulate the effector adenylate cyclase and thereby regulate numerous physiological processes such as cardiac function, cell proliferation, olfaction, and secretion (Muca and Vallar 1994, Mondry, Bourgeois et al. 1995, Asai, Yang et al. 1999, Geng, Ishikawa et al. 1999, Weinstein, Yu et al. 2001, Lohse, Engelhardt et al. 2003, Wettschureck and Offermanns 2005, Plagge, Kelsey et al. 2008, Layden, Newman et al. 2010, Mamillapalli and Wysolmerski 2010, Syrovatkina, Alegre et al. 2016, Cong, Xu et al. 2019). As all Gα subunits, Gα_s_ inactivates itself by the intrinsic hydrolysis of the bound GTP (GTPase) – a process that determines the duration of G protein-mediated signaling (Sprang 1997). The GTPase rates of most Gα subunits can be measured and compared *in vitro* using purified proteins. Specifically, Gα_s_ was reported to have a higher intrinsic GTP hydrolysis rate compared to other Gα subunits, ∼1 min^-1^ at 4°C (Krumins and Gilman 2002). The GTPase rate of G_i_ subfamily members was 0.1-0.2 min^-1^ or lower in the same conditions, and that of the G_12/13_ subfamily was even lower, at higher temperatures (Krumins and Gilman 2002). Furthermore, the GTPase activity of membersof the G_i_ and G_q_ subfamilies is accelerated by “canonical” RGS proteins, which act as GTPase activating proteins (GAPs) via a ∼120 amino acid “RGS domain” contained in all of these RGS proteins (Berman, Kozasa et al. 1996, Hunt, Fields et al. 1996, Koelle and Horvitz 1996, Siderovski, Hessel et al. 1996, Watson, Linder et al. 1996). The G_12/13_ subfamily can be inactivated by “non-canonical” RGS proteins from the RhoGEF family, which can exhibit GAP activity towards these particular Gα subunits (Sprang, Chen et al. 2007). In contrast, numerous studies have shown that none of the 20 canonical RGSs have any GAP activity towards the G_s_ subfamily (Berman, Kozasa et al. 1996, Hunt, Fields et al. 1996, Natochin and Artemyev 1998, Masuho, Balaji et al. 2020). One previous study suggested that a non-canonical RGS, RGS-PX1, which contains a divergent RGS domain, accelerated the GTPase activity of purified Gα_s_ and modulated Gα_s_ activity in cells (Zheng, Ma et al. 2001). However, no follow-up studies have been able to validate this suggestion and it is generally considered that no GAP for Gα_s_ has been identified to date (Siderovski and Willard 2005, Xie and Palmer 2007, Tesmer 2009, Sjogren, Blazer et al. 2010). It is therefore unclear what controls the inactivation rate of the G_s_ subfamily in a physiological context, and ways to investigate this *in vivo* by directly modifying the inactivation of Gα_s_ are lacking.

Numerous biochemical and structural studies have shown that inactivation of Gα subunits from the G_i/q_ subfamilies by canonical RGS domains involve interactions with both of the domains of the Gα subunit – the Gα “GTPase domain” and the “helical domain”. Many studies have suggested that interactions with the Gα GTPase domain are central to mediating RGS GAP activity (Tesmer, Berman et al. 1997, Milligan and Kostenis 2006, Soundararajan, Willard et al. 2008, Kosloff, Travis et al. 2011, Nance, Kreutz et al. 2013, Asli, Sadiya et al. 2018, Asli, Higazy-Mreih et al. 2021). This domain contains three flexible regions designated switch I, II and III that change conformation upon nucleotide binding and hydrolysis (Gilman 1995, Sprang, Chen et al. 2007). Previous work from our lab showed that conserved interactions of RGS residues with the Gα switch I and II regions of G_i_ subfamilies members are crucial for RGS activity, and quantified the reciprocal effects of non-conserved RGS “modulatory” residues that enhance interactions (Kosloff, Travis et al. 2011, Asli, Higazy-Mreih et al. 2021). In contrast, a third class of RGS “disruptor” residues were shown to attenuate interactions with specific Gα subunits, mostly via interactions with the Gα helical domain (Asli, Sadiya et al. 2018, Asli, Higazy-Mreih et al. 2021). Relevantly, the switch regions of Gα_s_ diverge relative to the G_i/q_ subfamily members that do interact with RGS proteins, and such differences might underlie the inability of RGS domains to interact with and inactivate Gα_s_ (Tesmer, Berman et al. 1997). A previous study implicated a Gα_s_-specific aspartate residue in switch II (D229) as an important contributor to the inability of Gα_s_ to interact with canonical RGS domains (Natochin and Artemyev 1998). Substitution of Gα_s_ D229 to serine, the corresponding residue in Gα_i1_, enabled some RGS16 GAP activity towards this Gα_s_ mutant. A following study showed that the reciprocal S202D mutation in Gα_t_ (a G_i_ subfamily member) severely impaired RGS16 GAP activity (Natochin and Artemyev 1998). However, the Gα_s_ D229S substitution resulted only in a small gain of function; RGS16 was reported to accelerate the GTPase rate of this mutant by only ∼5-fold (Natochin and Artemyev 1998), compared to a ∼1000-fold acceleration towards members of the G_i_ subfamily (Ross and Wilkie 2000).

On the other hand, previous studies pointed to the Gα helical domain as a central element that can attenuate interactions with RGS proteins in a specific way. Structural studies observed that Gα helical domain contacts with RGS domains can be heterogenic, suggesting this domain might contribute to interaction specificity towards different RGS proteins (Slep, Kercher et al. 2008, Soundararajan, Willard et al. 2008, Kimple, Soundararajan et al. 2009, Nance, Kreutz et al. 2013, Taylor, Bommarito et al. 2016). Indeed, previous studies from our lab showed that “disruptor residues” in the RGS R12 subfamily contribute to interaction specificity by reducing GAP activity towards Gα_o_, and that this effect is mediated via specific and unfavorable interactions with the Gα helical domain (Asli, Sadiya et al. 2018). Subsequent studies showed that such specific attenuation of RGS GAP activity towards members of the G_i_ subfamily can also occur in the RGS RZ, R4, and R7 subfamilies, in all cases mediated by disruptive interactions with the Gα helical domain (Salem-Mansour, Asli et al. 2018, Israeli, Asli et al. 2019, Asli, Higazy-Mreih et al. 2021). The Gα helical domain was also shown to be the major determinant of the exquisite specificity of RGS2 towards Gα_q_, preventing interactions with the G_i_ subfamily (Kasom, Gharra et al. 2018). Taken together, specific RGS interactions with the Gα helical domain seem to attenuate GAP activity selectively across many Gα subunits. Relevantly, Gα_s_ has a divergent helical domain that contains insertions not present in any other Gα subfamily (Sunahara, Tesmer et al. 1997), which might interfere with RGS binding. Overall, it is unclear which of these unique Gα_s_ structural features underlie its inability to interact with RGS domains and prevent RGS GAP activity towards this protein.

Here, we use Gα_s_ as a representative of the G_s_ subfamily to understand what determines its inability to interact and be turned off by RGS domains. We combine a comparative structural analysis using AlphaFold2 models of the entire Gα_s_ protein with biochemical assays that measure GTPase activities to identify Gα_s_ structural elements that may underlie its inability to interact with RGS proteins and function as “disruptor elements”. We test our computational predictions using amino acid substitutions between Gα_i1_ and Gα_s_, leading to Gα_s_ mutants that can be inactivated by canonical RGS proteins just as efficiently as Gα_i1_. These Gα_s_ mutants can be used to enable Gα_s_ inactivation by RGS proteins and thereby investigate the physiological consequences of manipulating Gα_s_ inactivation in cells.

## Results

### RGS16 does not accelerates the GTPase activity of Gα_s_

We compared the ability of RGS16, as a representative for high-activity RGS domains (Asli, Higazy-Mreih et al. 2021), to inactivate Gα_s_ and Gα_i1_ using single turnover GTPase assays. The average basal GTPase rate of Gα_s_ was 1.2 ± 0.2 min^-1^ (Fig. 1A), while the basal GTPase rate of Gα_i1_ was 4-fold lower, 0.3 ± 0.05 min^-1^ (Fig. 1B). As a control, we also measured under the same conditions the basal GTPase activity of Gα_o_, a related G_i_ subfamily member, which was 0.2 ± 0.01 min^-1^. As expected from previous studies (Krumins and Gilman 2002), RGS16 had no noticeable effect on GTP hydrolysis by Gα_s_ in the presence of 5-fold molar excess (2 μM) of RGS16 (Fig. 1A). In contrast, the GTPase rate of Gα_i1_ was accelerated to 2.2 min^-1^ in the presence of only 20 nM RGS16 (Fig. 1B). Dose response analysis with a larger concentration range of RGS16 confirmed that this RGS had no GAP activity towards Gα_s_, even at almost 100-fold molar excess of RGS16 relative to Gα_s_ (Fig. 1C). As expected, RGS16 showed very high GAP activity towards Gα_i1_, with an EC_50_ of 4 nM (Fig. 1C).

**Figure 1:**
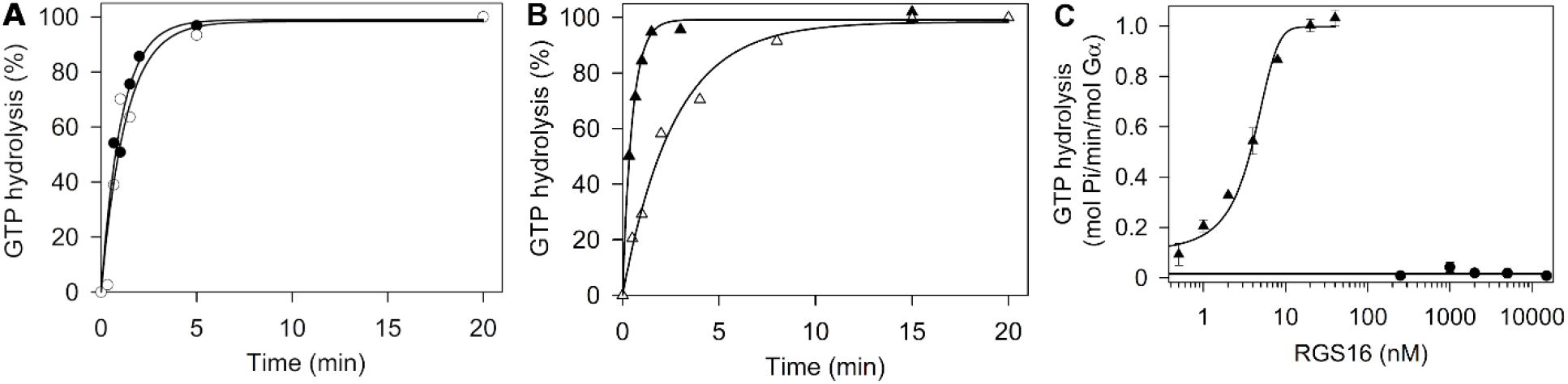
Gα_s_ GTPase activity is not affected by RGS16. (**A**) Representative single-turnover GTPase assays of Gα_s_ (400 nM), without RGS16 (open circles) and with 2 μM RGS16 (black circles). The average (n=3) reaction rate constant (k) was 1 ± 0.05 min^-1^ for Gα_s_ with RGS16 and 1.2 ± 0.2 min^-1^ for the basal activity of Gα_s_. **(B)** Representative single-turnover GTPase assays of Gα_i1_ (400 nM), with 20 nM RGS16 (black triangles) and without added RGS16 (open triangles). The average (n=3) reaction rate constant (k) was 2.2 ± 0.2 min^-1^ with RGS16 and 0.3 ± 0.05 min^-1^ for the basal activity of Gα_i1_. **(C)** Dose response analysis of RGS16 activity towards wild-type Gα_s_ (black circles) and wild-type Gα_i1_ (black triangles, EC_50_ = 4 ± 1 nM). Data presented are mean ± s.e.m. of experiments performed in triplicate, representative of three independent biological replicates each. Reaction rate constants for single-turnover GTPase assays were calculated using a single-exponential fit to the data using SigmaPlot 10.0, while EC_50_ values for dose response analysis of the net RGS-induced GTPase activity were calculated using three parameter sigmoidal curves.

### Gα_s_ contains unique structural elements that may determine its inability to interact with RGS domains

To identify putative Gα_s_ disruptor structural elements that may attenuate its interactions with RGS proteins, we compared Gα_s_ to experimental structures of Gα subunits from the G_i_ and G_q_ subfamilies in complex with RGS domains, the structural unit in RGS proteins that mediates all interactions with the Gα subunits. Notably, all available crystallographic structures of Gα_s_ are missing an insertion of ∼20 amino-acids – residues N66-D85, which are located in the α1-αA region of the Gα_s_ helical domain. We therefore used Alphafold2 to model the entire Gα_s_ subunit and compared it to complexes of Gα subunits in complex with RGS domains (Fig. 2). Relative to Gα_i1_, Gα_o_, and Gα_q_, Gα_s_ contains seven insertions: N66-D85 (α1-αA region), V114-P115, V134-D141 (the αB-αC loop), W277-S286, V301-K305, N315-D331 and S249-Y358 (Fig. 2A). The first three of these insertions are located in the Gα_s_ helical domain while the following four insertions are located in the GTPase domain. Alphafold2 produces five models for each protein input, and in the case of Gα_s_, all five models were generally similar except for the N66-D85 insertion; this region showed a variable conformation among the models (Fig. 2A), suggesting an inherent flexibility that might be the reason these regions were not resolved in any Gα_s_ experimental structure. Modeling a putative Gα_s_-RGS16 complex showed that all four GTPase domain insertions and the V114-P115 helical domain insertion are far away from the interface that has been characterized for other Gα subunits with RGS domains (Fig. 2A) – suggesting that these insertions cannot account for the inability of Gα_s_ to interact with RGSs. On the other hand, the Gα_s_ α1-αA region and the αB-αC loop are located close to the Gα interface with RGS domains observed in other Gα subunits and might introduce a steric clash preventing interactions with RGS proteins. In contrast, the corresponding two regions in Gα_i1_ are smaller and obviously do not clash with RGS domains (Fig. 2B). This raises the possibility that the Gα_s_ helical domain insertions in the α1-αA region or in the αB-αC loop might perturb binding to RGS domains by functioning as disruptor elements.

**Figure 2:**
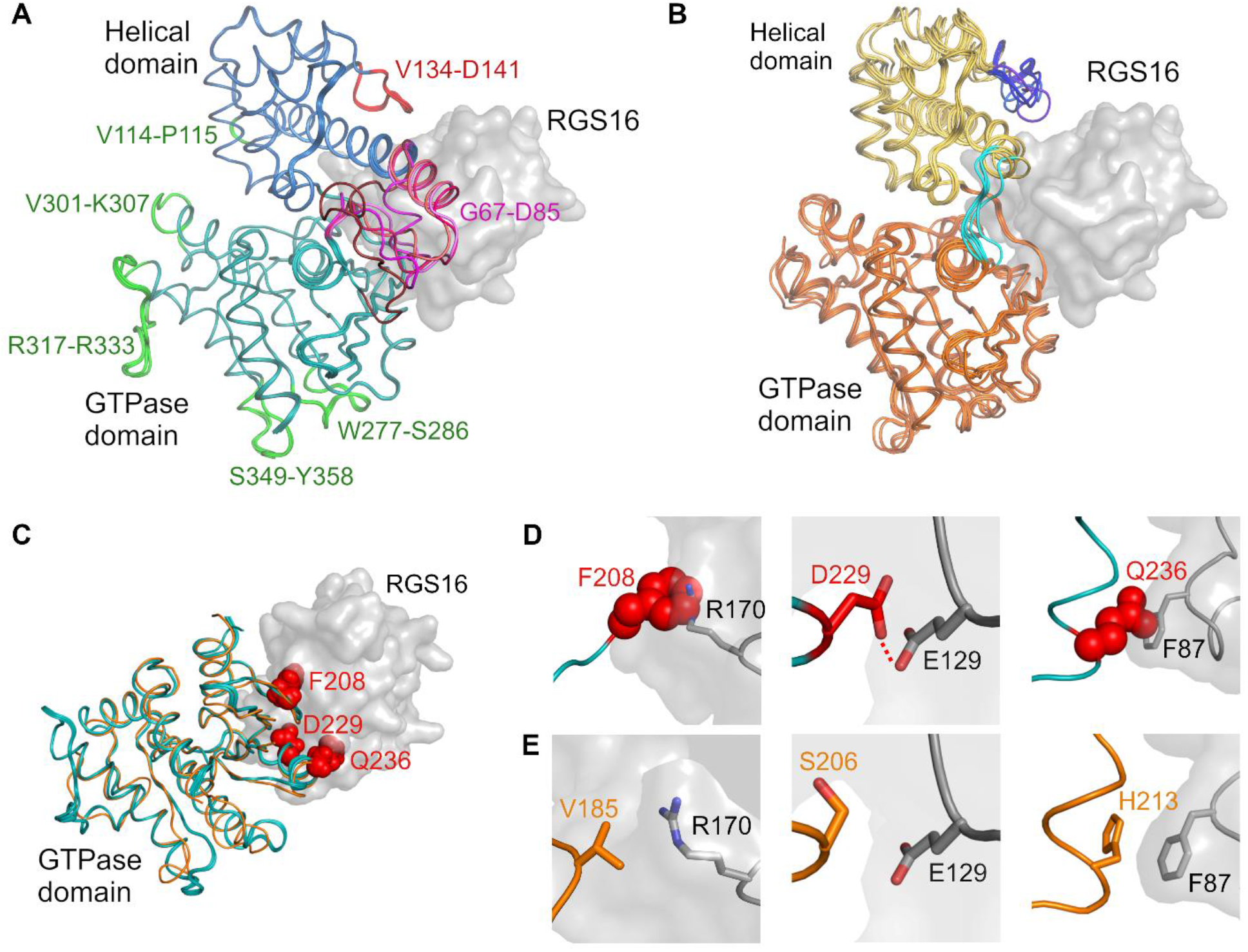
Structural elements in Gα_s_ that may disrupt interactions with RGS domains. (**A**) Alphafold2 models of wild-type Gα_s_, modelled bound to RGS16. The five models of Gα_s_ produced by Alphafold2 are shown in tube representation, colored teal (GTPase domain) and blue (helical domain). Five Gα_s_-unique insertions, four in the GTPase domain and one in the helical domain, that are far away from the RGS domain, are colored green. Two Gα_s_ regions that contain Gα_s_-unique insertions that are close to the putative location of the RGS domain and may introduce a steric clash are the αB-αC loop (V134-D141, red) and the α1-αA region (G67-D85, shades of magenta). The α1-αA region showed a variable conformation among the Alphafold2 models. **(B)** Representative experimental structures of Gα subunits that interact with high-activity RGS proteins (Gα_i1_, Gα_o_, Gα_q_). Gα subunits from five representative crystal structures (PDB IDs 2GTP, 1AGR, 2IK8, 3C7K, 4EKD) were superimposed, with Gα subunits shown as tubes colored orange (GTPase domain) and gold (helical domain). Gα αB-αC loops are colored in shades of blue and the α1-αA regions are colored cyan. In both panels, RGS16 from the Gα_i1_–RGS16 complex (2IK8) is shown as a transparent gray molecular surface. **(C)** Model of the Gα_s_ GTPase domain with RGS16. Gα_s_ from PDB ID 1AZS and Gα_i1_ with RGS16 from PDB ID 2IK8 were superimposed. Three Gα_s_ residues that may disrupt interaction with RGS domain are shown as red spheres. Gα_s_ and Gα_i1_ were visualized as in A and B, respectively, rotated 30° about the X-axis relative to A and with the Gα helical domain omitted for clarity. **(D)** Close up view of the predicted unfavorable interactions of the three putative Gα_s_ “disruptor residues” shown in C (red spheres or sticks) with RGS16 residues (gray sticks). A predicted unfavorable electrostatic interaction between Gα_s_ D229 and RGS16 E129 is marked with a dashed red line. **(E)** The corresponding Gα_i1_ residues, in relation to the same RGS16 residues shown in D.

We also searched for putative disruptor residues in the Gα_s_ GTPase domain by comparing the Gα_s_ sequence to our previous analysis using energy calculations of G_i_ and G_q_ interactions with RGS domains (Kasom, Gharra et al. 2018, Asli, Higazy-Mreih et al. 2021). Looking for residues that are unique to Gα_s_ and are located in positions that were shown to contribute to Gα_i/q_ interactions with RGSs, we identified three such Gα_s_ positions – F208, D229 and Q236. Gα_s_ residues F208 and Q236 are predicted to introduce distinct steric clashes with separate regions of RGS16 (Fig. 2D), while the negative charge of Gα_s_ D229 is predicted to repulse RGS16 E129 in a third RGS region – thereby pertubing interactions with the RGS domain (Fig. 2D). In contrast, the corresponding residues in Gα_i1_ (V185, S206, and H213) interact favorably with residues in the RGS domain (Fig. 2E). We note that each of these putative disruptor Gα_s_ residues is adjacent to a different region in the RGS16-Gα_i_ complex – F208 clashed with RGS16 helix α8, D229 with the RGS16 α5-α6 loop, and Q236 with the RGS16 α3-α4 loop (Asli, Higazy-Mreih et al. 2021). These Gα_s_ GTPase domain residues can thereby also perturb binding to RGS domains and function as disruptor elements.

### Substitution of the putative GTPase-domain Ga_s_ disruptor residues with the corresponding Gα_i1_ residues in the Gα_i1_ subunit abolish RGS16 activity towards Gα_i1_

To test whether the three residues in the Gα_s_ GTPase domain (F208, D229, and Q236), function as disruptor residues, we substituted the corresponding Gα_i1_ residues with their Gα_s_ counterparts (Fig. 3). These substitutions did not reduce Gα basal GTPase activity but did reduce the GAP activity of RGS16. The single Gα_i1_ mutants V185F and H213Q had either no effect (V185F) or a small effect (H213Q) on RGS16 GAP activity, respectively. The double mutant (V185F and H213Q) had a moderate effect on RGS16 GAP activity, with an EC_50_ of 45 ± 5, compared to an EC_50_ of 4 ± 1 with wild-type Gα_i1_. On the other hand, RGS16 GAP activity was abolished in both the single Gα_i1_ S206D mutant and the triple Gα_i1_ mutant (V185F, S206D, and H213Q).

**Figure 3:**
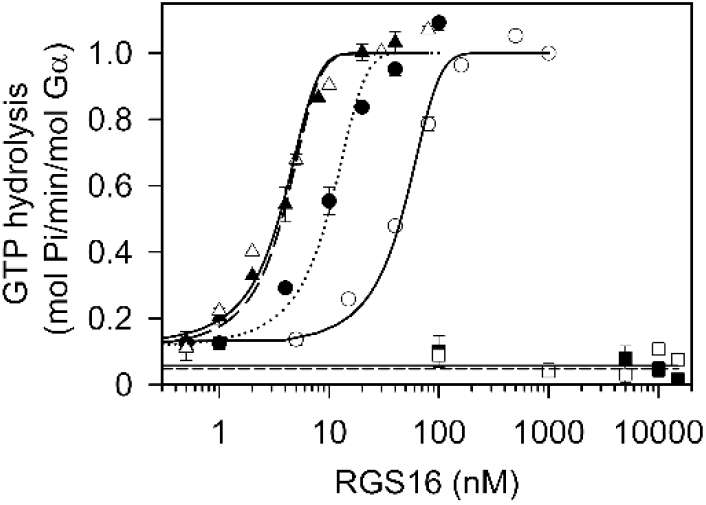
Substituting Gα_i1_ GTPase domain residues with the corresponding Gα_s_ putative disruptor residues is sufficient to abolish RGS16 GAP activity. Dose-response analysis of RGS16 activity toward wild-type Gα_i1_ (black triangles), Gα_i1_-V185F single mutant (open triangles and dashed line), Gα_i1_-S206D single mutant (open squares), Gα_i1_-H213Q single mutant (black circles), Gα_i1_-V185F/H213Q double mutant (open circles), and Gα_i1_-V185F/S206D/H213Q triple mutant (black squares). EC_50_ values, calculated as in Fig. 1C, are as follows: wild-type Gα_i1_ = 4±1 nM, Gα_i1_-V185F single mutant = 4±1 nM, Gα_i1_-H213Q single mutant = 7±1 nM, and Gα_i1_-V185F/H213Q double mutant = 45±5 nM. Data presented are mean ± s.e.m. of experiments performed in triplicate, representative of at least three independent biological replicates each.

### The putative GTPase-domain Gα_s_ disruptor residues are necessary and sufficient to prevent RGS16 GAP activity

We next performed the reciprocal experiment, substituting the putative Gα_s_ GTPase domain disruptor residues with the corresponding residues from Gα_i1_. Substituting Gα_s_ D229 with serine enabled RGS16 GAP activity (Fig. 4), in agreement with previous results (Natochin and Artemyev 1998). However, this GAP activity (EC_50_ = 800 ± 200) is more than two orders of magnitudes lower than the GAP activity we measured towards Gα_i1_ (EC_50_ = 4 ±1, Fig. 1C). Substitution only of Gα_s_ F208 and Q236 with valine and histidine did not enable any RGS16 GAP activity (Fig. 4). On the other hand, substituting all three putative Gα_s_ disruptor residues simultaneously with the corresponding Gα_i1_ residues led to full gain-of-function, enabling high RGS GAP activity towards Gα_s_ that is indistinguishable from the high RGS GAP activity towards Gα_i1_ (Fig. 4 cf. Fig. 1C).

**Figure 4:**
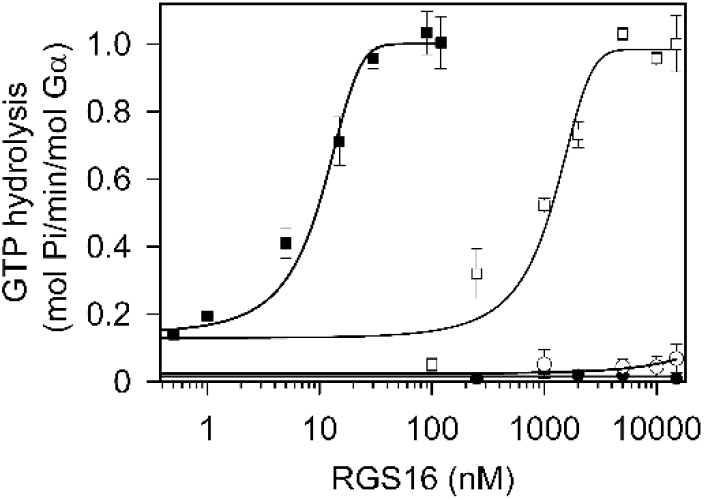
Substituting all three putative Gα_s_ disruptor residues with the corresponding Gα_i1_ residues results in full RGS16 gain of function. Dose-response analysis of RGS16 activity toward wild-type Gα_s_ (black circles), Gα_s_-D229S single mutant (open squares), Gα_s_-F208V/Q236H double mutant (open circles), and Gα_s_-F208V/D229S/Q236H triple mutant (black squares). EC_50_ values, calculated as in Fig. 1C, are as follows: Gα_s_ D229S single mutant = 800±200 nM, and Gα_s_ F208V/D229S/Q236H triple mutant = 9±2 nM. Data presented are mean ± s.e.m. of experiments performed in triplicate, representative of at least three independent biological replicates each.

## Discussion

Our results show that three residues in the Gα_s_ GTPase domain are responsible for preventing interactions with RGS proteins and function together as disruptor elements – a phenylalanine (F208) in switch I and two residues in switch II, D229 and Q236. Each of these three disruptor Gα_s_ residues interacts with a different region of the RGS protein. These three Gα_s_ residues are both necessary and sufficient to explain the inability of RGS proteins to inactivate members of the Gα_s_ subfamily. On the other hand, our results show that the helical domain of Gα_s_, despite the unique and large insertions it contains, does not introduce a steric clash that can attenuate interactions with RGS domains. Quantifying the effect of the three Gα_s_ disruptor residues, D229 had the stronger effect, preventing any RGS GAP activity, consistent with (Natochin and Artemyev 1998), while the two other residues, F208 and Q236, attenuate RGS GAP activity. We note a Gα mutant that correspond to the S206D single mutation is sufficient to block RGS GAP activity towards the G_i_ and G_q_ subfamilies (Fig. 3) – similar to the well-studied G183S mutation in Gα_i1_, which has been previously shown to prevent interactions with RGS proteins (Lan, Sarvazyan et al. 1998, Fu, Zhong et al. 2004). However, to enable full RGS GAP activity towards Gα_s_, as high as the GAP activity toward Gα_i1_, all three Gα_s_ disruptor residues needed to be substituted with their Gα_i1_ counterparts.

We note that in previous studies, we showed that in the G_i_ subfamily, attenuation of specific RGS proteins was mediated mostly by interactions with the Gα helical domain (Asli, Sadiya et al. 2018, Kasom, Gharra et al. 2018, Asli, Higazy-Mreih et al. 2021). In contrast, evolution placed all of the Gα_s_ disruptor elements in the GTPase domain, which is the central domain that mediates Gα interactions with diverse partners (Navot and Kosloff 2019), so in Gα_s,_ the helical domain does not play a role in preventing interactions with RGS proteins.

Interestingly, some RGS proteins having low activity towards the G_i_ subfamily contain disruptor residues that interact with the same Gα structural regions (Asli, Higazy-Mreih et al. 2021), suggesting evolution has used disruptor elements in reciprocal structural locations in either the Gα or the RGS side of the interface to achieve interaction specificity at the family level.

Furthermore, the fact that the activity of the G_s_ subfamily is not downregulated by any RGS protein highlights a central unanswered question – what does control the inactivation rate of Gα_s_ beside its intrinsic GTPase activity *in vivo*? Relevantly, our results show that Gα_s_ has a higher intrinsic GTPase activity compared to the G_i_ subfamily, which corroborates measurements performed previously (Krumins and Gilman 2002). However, whether this higher Gα_s_ intrinsic GTPase is sufficient to regulate its function *in vivo* is still unclear. Therefore, the Gα_s_ mutants we identified here can be used to investigate this question by modifying its inactivation *in vivo*.

## Acknowledgements

This work was supported by the Israel Science Foundation (grant 1454/13) to MK, grants from the Canadian Institutes of Health Research (CIHR: PJT-183758) and the Natural Science and Engineering Research Council of Canada (NSRC:RGPIN-2019-05556) to MB, the International Development Research Centre (IDRC), the Israel Science Foundation (ISF), and the Azrieli Foundation (grant 3512/19) to MK and MB, and by a grant from the Council for Higher Education through the Data Science Research Center at the University of Haifa to MK.

## Materials and methods

### Protein sequences, structures and models

We used the following 3D structures in our analysis and visualization of Gα-RGS complexes (with PDB codes for each structure): Gα_i1_–RGS16 (2IK8) (Soundararajan, Willard et al. 2008), Gα_i1_–RGS4 (1AGR) (Tesmer, Berman et al. 1997), Gα_i1_–RGS1 (2GTP) (Soundararajan, Willard et al. 2008), Gα_o_–RGS16 (3C7K) (Slep, Kercher et al. 2008), and Gα_q_–RGS2 (4EKD) (Nance, Kreutz et al. 2013). The structure of full length Gα_s_ was predicted using the Alphafold2 ColabFold server (https://colab.research.google.com/github/sokrypton/ColabFold/blob/main/AlphaFold2.ipynb). 3D structural visualization and superimposition were carried out with the molecular graphics program PyMol (http://pymol.org).

### Protein expression, purification and mutagenesis

All proteins were expressed as N-terminally His_6_-tagged proteins. The sequence of Gα_s_ used here corresponds to UNIPROT ID P63092-1 and that of Gα_i1_ to UNIPROT ID P63096. The RGS domain of human RGS16 (residue 86–205) and human Gα_i1_ (residues 31-354) were expressed in the pNIC-SGC1 vector (Addgene), and pProEXHTb vector (Invitrogen), respectively. Mutations were introduced using the QuikChange site-directed mutagenesis kit (Invitrogen). Expression and purification were conducted as described in (Kasom, Gharra et al. 2018). Human Gα_s_ (residues 36-393) was expressed in Escherichia coli BL21 (DE3) cells and grown in 1 L of TY broth at 30°C, until an A600 nm ≥ 1.4 was reached. The temperature was then reduced to 25°C and protein expression was induced by the addition 30 µM isopropyl-D-thiogalactopyranoside. After 16–18 h, cells were harvested by centrifugation at 6000 g for 30 min at 4°C, followed by freezing the pellets at −80°C. Gα subunits were purified as described in (Kasom, Gharra et al. 2018). The yield and purity of all proteins was validated by SDS-PAGE electrophoresis and Coomassie staining. Protein concentrations were determined by measuring absorption at A_280_ nm, using extinction coefficients predicted with ProtParam (Swiss Institute for Bioinformatics) with the sequence of each expressed protein.

### Single-turnover GTPase assays and RGS GAP activity measurements

We used single-turnover GTPase assays to measure the GTPase activity of Gα proteins with/without RGS proteins (Ross and Wilkie 2000, Kosloff, Travis et al. 2011). Gα subunits were loaded with radioactively labeled GTP by incubating for 20 min at 20°C (Gα_s_) and for 15 min at 30°C (Gα_i1_) with 1 μM [γ-^33^P]-GTP (Hartmann Analytic GmbH) or 1 μM [γ-^32^P]-GTP (Perkin Elmer) in Reaction Buffer (50 mM HEPES, pH 7.5, 0.05% polyoxyethylene (v/v), 5 mM EDTA, 5 mg/ml BSA, 1 mM dithiothreitol), followed by cooling on ice for 5 min. GTP hydrolysis was measured at 4°C, initiated by raising the free magnesium concentration to 5 mM for Gα_s_ and 1 mM for Gα_i1_ (using MgCl_2_), together with 100 μM cold GTP (final concentration), with or without added RGS proteins. Aliquots were taken at different time points and quenched with 5% charcoal in 50 mM Na_2_H_2_PO_4_ (pH 3), followed by centrifugation at 12,000 g for 5 min at room temperature. The supernatants were transferred to 3 ml liquid scintillation liquid and analyzed using a Tri-Carb 2810 TR scintillation counter (Perkin Elmer).

We used dose-response analysis with a range of RGS concentrations to measure GAP activity towards Gα subunits. Gα subunits were incubated with 1 μM [γ-^33^P or γ-^32^P]-GTP for 20 min at 20°C (Gα_s_) and for 15 min at 30°C (Gα_i1_) and then cooled on ice for 5 min. Hydrolysis was initiated by adding 10 μl RGS protein in different concentrations in assay buffer containing 1 mM MgCl_2_ for Gα_i1_ or 5 mM MgCl_2_ for Gα_s_, together with 100 μM cold GTP to a tube containing 20 μl Gα subunit (500 nM) on ice. Each reaction was terminated after 30 sec by adding 100 μl 5% perchloric acid and quenched with 700 μl 10% (w/v) charcoal slurry in 50 mM phosphate buffer (pH 7.5), followed by centrifugation at 12,000 g for 5 min at room temperature. 200 μl of the supernatants were transferred to 3 ml liquid scintillation and analyzed using a Tri-Carb 2810 TR scintillation counter (Perkin Elmer).

